# Influence of normobaric hypoxic exercise on endothelial progenitor cell senescence by *in vitro* cultivation with conditioned media

**DOI:** 10.1101/2020.02.16.949677

**Authors:** Jan-Frieder Harmsen, Dennis Nebe, Klara Brixius, Alexander Schenk, Wilhelm Bloch

## Abstract

**Objective:** Circulating endothelial progenitor cells (EPCs) were shown to be affected in cardiovascular and metabolic diseases. As interventional strategies, hypoxia and exercise are both known to increase the number and enhance the function of EPCs, potentially by extending their lifespan induced by a reduced senescence. Therefore, this pilot study investigated the effect of exercise under normobaric hypoxia on the senescence of EPCs by *in vitro* cultivation with autologous human serum (AHS).

**Methods:** Four healthy trained young males (23 ± 2 years) performed an incremental cycling step test until exhaustion in a normobaric hypoxic-chamber with an average altitude of 4,000 m (O_2_ 12.3%). Blood serum was taken at pre, 10 min post and 4 h post, which was later used for *in vitro* cultivation of EPCs. Senescence was investigated by ß-galactosidase staining.

**Results:** The participants spent 30-40 min in normobaric hypoxia. The EPC senescence rate was reduced 10 min (0.72 ± 0.57%) and 4 hours (0.67 ± 0.52%) after exercise compared to pre (1.89 ± 0.37%).

**Conclusion:** This pilot study indicates that intense exercise under normobaric hypoxia may enhance EPC function by slowing down their senescence.

## Introduction

Endothelial progenitor cells (EPCs) are described as CD34^+^ Flk-1^+^ possessing the ability to differentiate into endothelial cells *in vitro* and thereby integrate into sites of active angiogenesis [1,2]. Stimuli inducing the release of EPCs from bone marrow into the circulation include acute injury, tissue ischemia, tumor angiogenesis and myocardial infarction [3]. In the last two decades, EPCs became a promising target of regenerative medicine research and several interventional approaches have been performed to investigate the mobilization of EPCs. In particular, abundant research has been conducted showing enhanced levels of circulating EPCs after exercise interventions in patients [4–6] as well as in the healthy population [7,8].

Besides physical exercise, the exposition to normobaric hypoxia for a short time period (1 hour) was also shown to be sufficient to increase EPCs in the circulation [9]. Thus, combining exercise and hypoxia in an attempt to maximize the stimulus of oxygen deprivation has even been shown to be superior to normobaric exercise in terms of cardiac and muscular adaptations possibly by enhancing EPC mobilization [10]. For the purpose of determining the optimal exercise stimulus in hypoxia, Tsai and colleagues [7] identified high intensity interval training to stimulate EPC production more than high volume training.

Independent of an upregulation of EPC formation, physical exercise and/or hypoxia may enhance EPC count indirectly by extending their lifespan [11] represented by downregulated cell senescence factors. However, the direct effect of exercise under normobaric hypoxia specifically on the senescence of EPCs has not been studied to date. Therefore, the aim of this pilot study was to investigate the influence of acute exhausting exercise under normobaric hypoxia by *in vitro* cultivation of EPCs with autologous human serum.

## Methods and materials

### Participants

Four healthy male sport students participated in this study. Characteristics of basic anthropometric and physical parameters of the participants are summarized in Table 1. All participants abstained from alcohol consumption for 24 h before the intervention and were nonsmokers. All participants gave written informed consent to contribute to the study. The study has been performed according to the Declaration of Helsinki, and the local institutional review board has approved the procedures.

**Table 1.** Anthropometric data and physical parameters (n=4); Mean ± SD.

|  |  |
| --- | --- |
| Age [years] | 23 ± 2 |
| Height [cm] | 181 ± 6 |
| Body weight [kg] | 74.9 ± 8.7 |
| Training hours per week | 10.3 ± 3.7 |

### Exercise study protocol

Every subject performed an incremental cycling step test until exhaustion under normobaric hypoxia. Before the test started, participants performed a 10 min warm-up at 70 W and rested for 5 min afterwards. The intensity of the subsequent test started at 100 W and was increased every three minutes by 30 W. Participants had to maintain a pedal rate of 70-80 rpm. The test was aborted as soon as the pedal rate dropped below 60 rpm. A Cyclus 2 (RBM Elektronik, Automation, Leipzig, Germany) cycling ergometer was used for the incremental cycling test. The hypoxic condition was induced by using a normobaric hypoxic-chamber (Hypoxic Training Systems, Hypoxico, New York) with an average altitude of 4,000 m (O_2_ 12.3%). Further information can be obtained from Suhr and colleagues [12] as the same altitude chamber and system to determine maximal oxygen uptake (VO_2 peak_) was used.

### Blood sampling and serum preparation

Blood samples were taken by venipuncture before (pre-), 10 min post- and 4 h post-exercise. Blood was drawn into SST vacutainer (Becton Dickinson GmbH, Heidelberg, Germany) for serum preparation. Blood samples in SST vacutainers were stored at 7°C for 30 min for blood clotting and then centrifuged for 10 min at 1861xg and 4°C for serum preparation. The blood serum was aliquoted and stored at −80°C until use.

### Isolation of EPCs from whole blood

EDTA blood samples were taken from each participant for isolation of peripheral blood mononuclear cells (PBMCs) at rest. PBMCs were isolated by centrifugation for 30 min at 800xg with a lymphocyte separation medium (PAA Laboratories GmbH, Cölbe, Germany). EPC isolation was modified from Funke et al. 2008 [13]. In brief, PBMCs were cultivated in 24-well plates containing fibronectin-coated glass plates with MV2-endothelial cell basal medium (Promo Cell GmbH, Heidelberg, Germany) for 1 week. After 4 days medium was changed.

### EPC cultivation with conditioned medium

Isolated EPCs were cultivated with MV2-endothelial cell basal medium containing 5% human blood serum of all measurement time points (pre, 10 min post, 4 h post) for 24 h. As a control medium, 5% fetal bovine serum (FBS) was used.

### β-galactosidase staining

Cultivation medium was removed after 24 h of cultivation. EPCs were washed with PBS and fixated with a fixation solution containing PBS, 0,2% glutaraldehyde, 5mM EGTA (ethylene glycol-bis(β-aminoethyl ether)-N,N,N’,N’-tetraacetic acid) and 2mM MgCl_2_ for 15 min. After fixation EPCs were washed three times for 5 min with a wash solution containing PBS, 0,01% natrium-desoxycholate, 0,02% Nonident P40, 5mM EGTA and 2mM MgCl_2_. Following, staining was performed for 24 h in the dark at 37°C with a fresh prepared staining solution consisting of 10mM potassium ferricyanide (K_3_[Fe(CN)_6_]), 10mM potassium ferrocyanide (K_4_[Fe(CN)_6_]) and 1mg/ml X-gal (5-bromo-4-chloro-3-indolyl-β-D-galactopyranoside) added to the wash solution. After staining, cells were mounted with Vectashield (Vector Laboratories, Burlingame CA, USA) for conservation and simultaneous staining of nuclei with DAPI. EPCs were analyzed with a fluorescence microscope (Axiophot, Zeiss GmbH, Jena, Germany) using fluorescence for counting all cells and light microscopy for β-Gal positive cells on the same device.

## Statistics

Results are presented as means ± SD. Because of the small sample size, results are discussed in a descriptive manner instead of performing inferential statistics.

## Results

Participants had to abort the test after a duration ranging from 15 to 25 min resulting in a total time of 30-40 min spent in normobaric hypoxia. The VO_2 peak_ was 56.7 ± 7.7 ml · kg^-1^· min^-1^. The senescence rate was measured after *in vitro* cultivation with serum collected pre, 10 min and 4 hours after acute exercise under hypoxia. As shown in Figure 1, senescence was reduced 10 min (0.72 ± 0.57%) and 4 hours (0.67 ± 0.52%) after exercise compared to pre (1.89 ± 0.37%).

**Figure 1.**
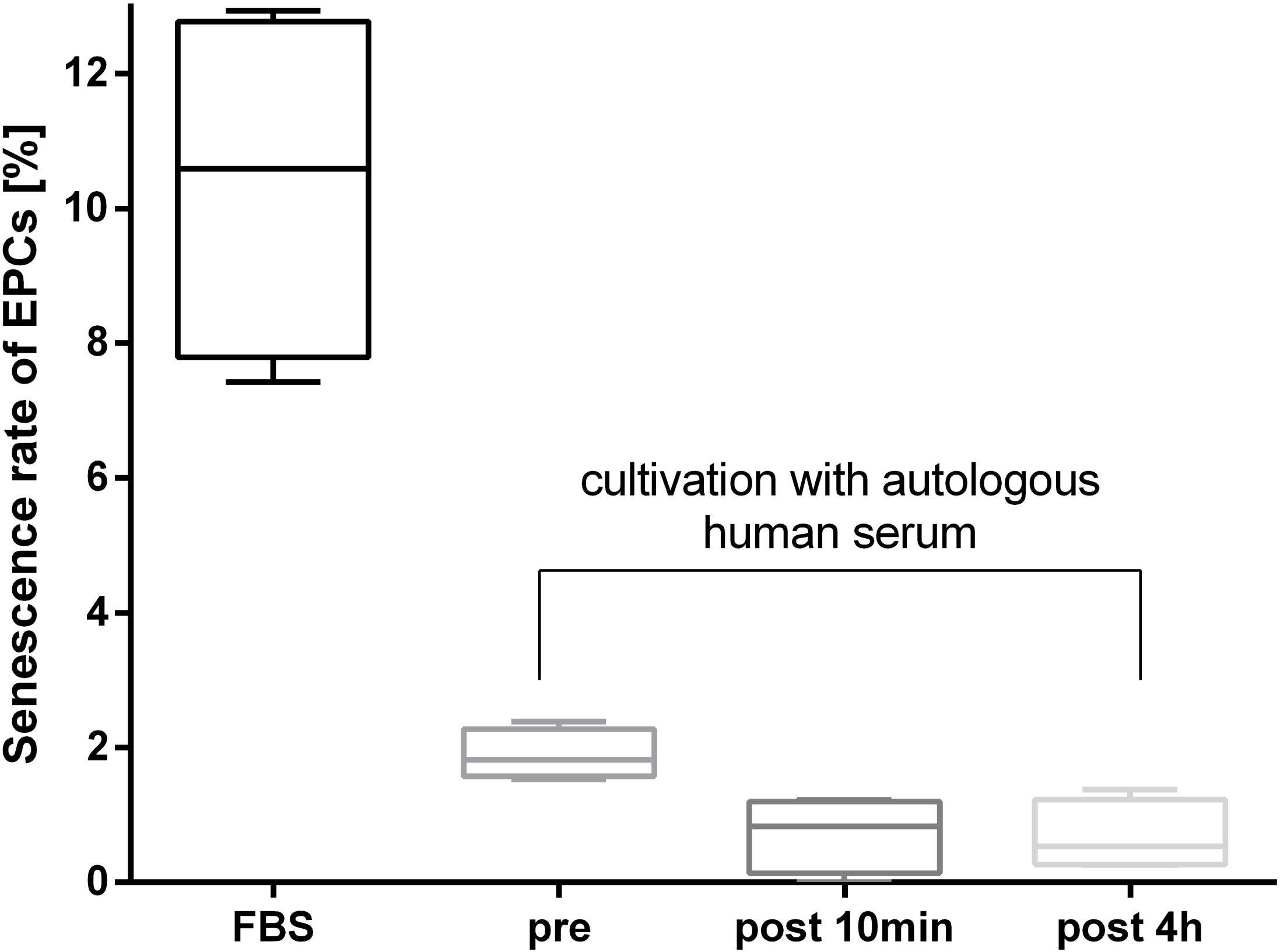
Senescence rate of endothelial progenitor cells (EPCs) after *in vitro* cultivation of EPCs with either fetal bovine serum (FBS) or autologous human serum (AHS) at pre, post 10min and post 4h.

In comparison to *in vitro* cultivation with FBS and AHS, cultivation with FBS led to a much greater senescence rate (10.38 ± 2.66%) than seen after all AHS conditions.

## Discussion

This pilot study indicates that *in vitro* cultivation of EPCs with conditioned media containing blood serum from exercise to exhaustion under normobaric hypoxia leads to a reduction in the senescence rate of EPCs. To the best of our knowledge, this is the first investigation demonstrating acute changes in EPC senescence in response to *in vitro* cultivation with blood serum before and after a single bout of intense hypoxic exercise. Investigating chronic adaptations, Wang et al. (2014) [10] showed an increase in the proliferative capacity of EPCs after five weeks of hypoxic exercise training at 60% VO_2 max_ leading to a larger improvement in aerobic capacity compared to a normoxic exercise control group. Taken together, these findings indicate that the conditioning of blood serum through exercise under normobaric hypoxia induces promising effects on the endothelial system. In this context, especially hypoxia is known to induce the release of cytokines like vascular endothelial growth factor (VEGF), erythropoietin (EPO) and hepatocyte growth factor (HGF) into the circulation [14]. Ceradini and Gurtner [14] found a direct correlation of EPC activation and serum EPO. In addition, HGF was consistently increased which has been shown to stimulate angiogenesis even more efficiently than VEGF [15]. Maximizing oxygen deprivation through exercise under hypoxia may generate the necessary blood serum milieu to slow down EPC senescence in the later *in vitro* cultivation.

A reduction of EPC senescence would potentially be a crucial outcome in a clinical setting contributing to a higher EPC count, which has been shown to be a positive prognostic marker in several pathologies. Among others, Werner and colleagues [16] demonstrated that differences in CD34+KDR+ EPCs could predict cardiovascular outcomes in patients with coronary artery disease. In addition, a range of putative EPCs has been shown to be lower and associated with long-term mortality in type 2 diabetes compared to healthy controls [17,18]. Nonetheless, our study highlights immense differences in the cultivation of EPCs with AHS in contrast to commercial FBS-supplementation. Senescence of primary human EPCs was explicitly accelerated after cultivation with FBS. It should be kept in mind, that incubation of primary human EPCs with FBS challenges the cells, i.a. promoting senescence, and therefore may overlie the effects of other stimuli that were tested. This finding is in line with a recent review of Bauman and colleagues [19] concluding that the sustained use of FBS leads to inconsistencies in cell propagation protocols impeding data comparison among different studies and laboratories and therefore deaccelerates the progress in EPC research.

When reviewing this research it is important to acknowledge that the sample size was very small (n=4) and consequently was not large enough to detect any effect with statistical power, therefore limiting the ability to draw firm conclusions. However, our data contribute to the growing body of evidence supporting the clinical benefits of hypoxic exercise [10,20]. As the characteristics of optimal training stimulus in hypoxia are still largely unclear, future trials with homogenous clinical populations need to address which exercise intensity at which altitude for how long is feasible, and concurrently sufficient to alter EPC behavior including e.g. their senescence.

## Acknowledgments

The results of the study are presented clearly, honestly, and without fabrication, falsification, or inappropriate data manipulation.

